# Duplications at 19q13.33 in patients with neurodevelopmental disorders

**DOI:** 10.1101/130377

**Authors:** Eduardo Pérez-Palma, Elmo Saarentaus, Joris Andrieux, Marie Ravoet, Giancarlo V. De Ferrari, Peter Nuernberg, Bertrand Isidor, Bernd A. Neubauer, Dennis Lal

## Abstract

**OBJECTIVE:** After recent publication of the first patients with disease associated missense variants in *GRIN2D*, we evaluate the effect of copy number variation (CNV) overlapping this gene towards the presentation of neurodevelopmental disorders.

**METHODS:** We explored ClinVar (N°CNV = 41,398) and DECIPHER (N°CNV = 30,222) clinical databases of genomic variations for patients with copy number changes overlapping the *GRIN2D* gene at the 19q13.33 locus and evaluated their respective phenotype alongside their frequency, gene content and expression with publicly available reference databases.

**RESULTS:** We identified 13 patients with microduplications at the 19q13.33 locus. The majority of CNVs arose *de novo* and comparable CNVs are not present in control databases. All patients were reported to have neurodevelopmental disorders and dysmorphic features as the most common clinical phenotype (N= 10/13), followed by seizures (N= 6/13) and intellectual disability (N= 5/13). All duplications shared a consensus region of 405 kb overlapping 13 genes. After screening for duplication tolerance in control populations, positive gene brain expression and gene dosage sensitivity analysis, we highlight four genes for future evaluation: *CARD8*, *C19orf68*, *KDELR1* and *GRIN2D,* which are promising candidates for disease causality. Further, investigation of the literature especially supports *GRIN2D* as the best candidate gene.

**CONCLUSIONS:** Our study presents dup19q13.33 as novel duplication syndrome locus associated with neurodevelopmental disorders. *CARD8*, *C19orf68*, *KDELR1* and *GRIN2D* are promising candidates for functional follow up.

## Introduction

N-methyl-D-aspartate (NMDA) receptors are involved in neurodevelopmental processes such as synaptogenesis, learning and memory. Structurally, NMDA receptors are composed by two subunits of GluN1 and GluN2 which are specifically encoded by the *GRIN1* and *GRIN2A* to *GRIN2D* genes, respectively^1^. While single nucleotide and Copy Number Variants (CNV) in the NMDA receptor subunits *GRIN1*, *GRIN2A* and *GRIN2B* have been associated with a range of neurodevelopmental disorders (NDDs), little is known about the association of *GRIN2D* variants and NDDs. Recently, *de novo* missense mutations in *GRIN2D* (p.Val667Ile) were identified as the cause of severe epileptic encephalopathy^2^ in two independent patients. However, whether CNVs covering the *GRIN2D* locus are also associated with disease has not been studied. *GRIN2D* is encoded at the end of the long arm of chromosome 19 (19q13.33) locus. We hypothesize that dosage changes in *GRIN2D* are highly likely to be disease associated based on the high sequence homology, expression during neurodevelopment and a functional relationship with the established disease associated paralogs genes.

## Methods

Using the gene oriented query “*GRIN2D*”, we accessed two publically available repositories of clinical genetic variation: 1) The Database of Chromosomal Imbalance and Phenotype in Humans using Ensembl Resources, DECIPHER^3^ (URL: https://decipher.sanger.ac.uk, accessed on July 2016), and 2) The public archive of interpretations of clinically relevant variants, ClinVar^4^ (URL: http://www.ncbi.nlm.nih.gov/clinvar, accessed on July 2016). For each of the identified patients, we contacted the individual scientist who deposited the entry to aquire further phenotype information including the presence of: Intellectual disability (ID), developmental delay (DD), seizures, hypotonia, dysmorphism (Dysm), learning difficulties, behavioral problems as well as social communication and behavioral disorders of the autism spectrum (ASD)^5^. All phenotypes evaluated were considered as a binary denominators (i.e. YES/NO). Gene annotations of the extracted CNVs refer to the genome build GRCh37/hg19. Consensus region was determined with an in-house python script (available upon request). Genes inside the consensus region were further evaluated as disease candidate genes with additional publicly available resources for: 1) Brain expression, strongly brain-expressed genes (n = 4,756), specified by a log (RPKM) >4.5 of the BrainSpan RNA.Seq transcriptome dataset^6^; 2) Overlapping CNVs reported in the curated control inclusive map of the Database of Genomic Variants^7^; 3) Loss of Function (LoF) Intolerance reported in the Exome Agregation consortium^8^, given by a probability of being LoF intolerant (pLI score) equal or greater than 0.9 based on the observed genetic variation of 60,706 healthy individuals; and 4) overlapping CNVs reported in 20,227 controls^9^. Genome wide brain-specific non-coding functional elements were extracted from the Genoskyline project (http://genocanyon.med.yale.edu/GenoSkyline) which implements a statistical framework based on high throughput genetic and epigenetic data in order to predict tissue specific functional non-coding elements^10^.

## Results

We detected 13 patients with CNVs overlapping the 19q13.33 locus (Table 1). Interestingly, all of them were duplications. Three were annotated in ClinVar (patients 1 through 3) and ten in DECIPHER (patients 4 through 13). Although a fourteenth individual did fulfill the criteria (DECIPHER entry 275388), given the actual size of the reported variant in comparison to the entire chromosome 19 (CNV = 58.83 Mb vs Chr19= 59.12 Mb), it was considered a chromosome trisomy and therefore excluded. Detailed clinical phenotypes are provided in Table 1. Notably, all patients were reported to have mild to severe forms of NDDs. Of all the phenotypes evaluated, mild but distinct dysmorphic features was the most frequent (n = 10), followed by seizures (n = 6, including generalized tonic and febrile seizures), developmental delay (n = 6) and intellectual disability (n = 5). In particular, dysmorphisms were present in patients carrying a CNV larger than 3mb (pathogenic size according to the American College of Human Genetics). The image of one of such patients is depicted in figure 1A, showing a child with signs of macrostomia, mid-face hypoplasia, and progenia.

**Table 1.**
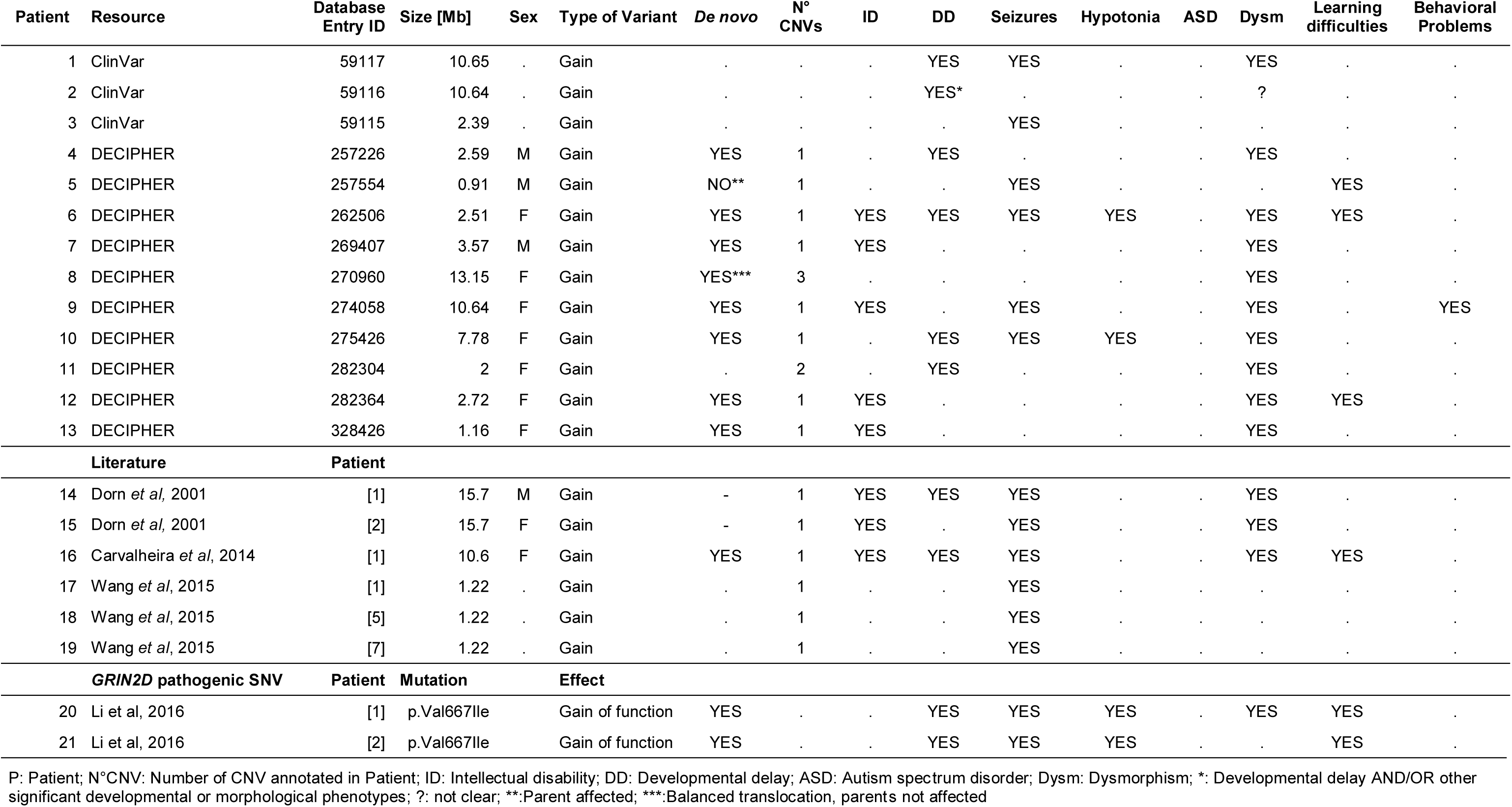
Clinical phenotypes of the 13 retrieved patients with GRIN2D variants at the 19q13.33 locus.

**Figure 1.**
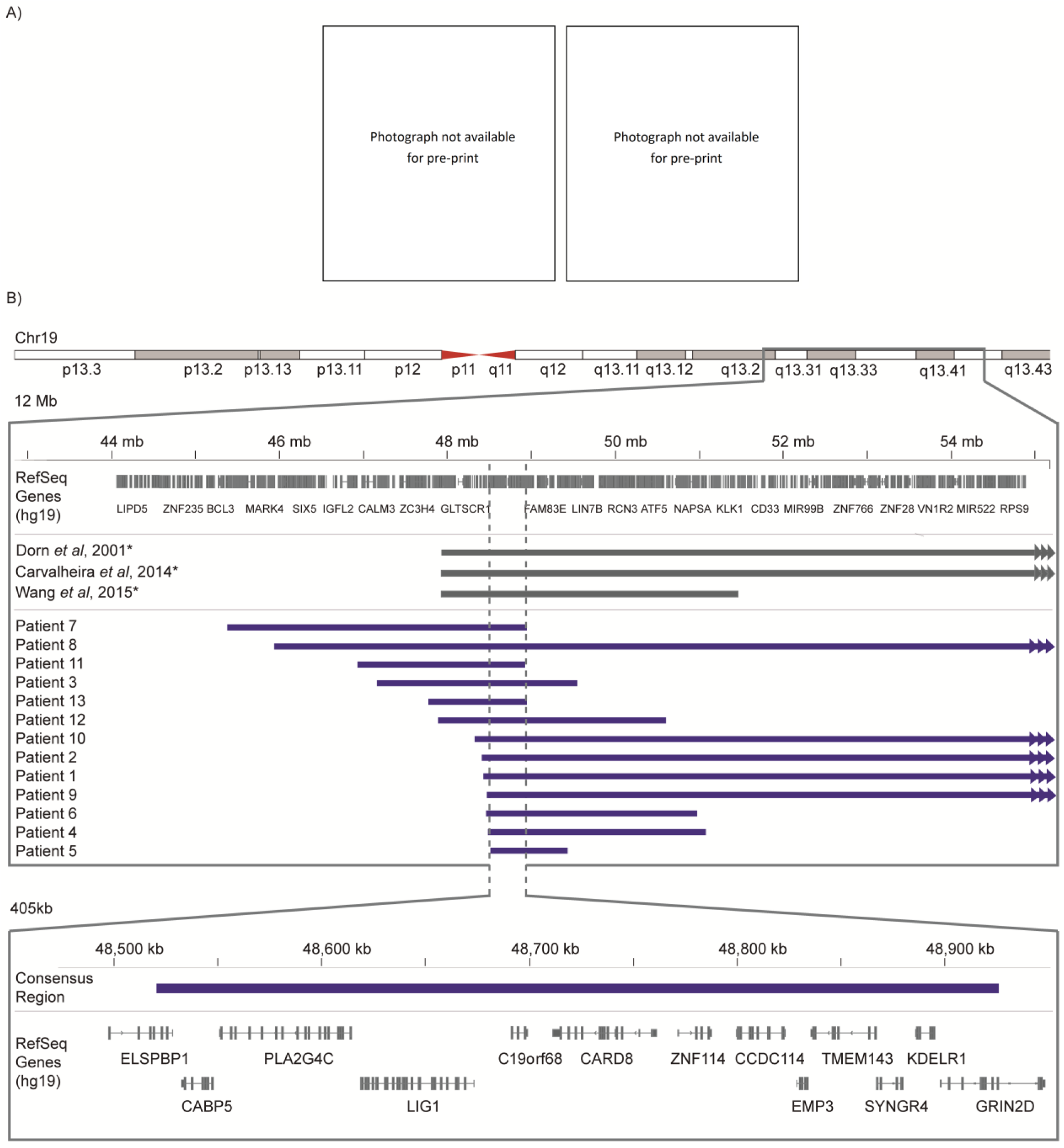
Genomic and facial overview of the micro duplications overlapping GRIN2D gene found in the retrieved patients. A) Clinical anteroposterior facial photograph of patient 12 depicting characteristic facial features. B) Upper panel. Thirteen patients were identified with GRIN2D duplications at the 19q13.33 locus. Blue horizontal bars represent the respective microduplication size and breakpoints according to GRCh37/hg19 human genome reference in a 12mb genomic window. Gray horizontal bars represent the respective microduplication reported in Dorn et al 2001, Carvalheira et al 2014 and Wang et al 2015 were no exact CNV boundaries are specified (*). Microduplications larger than the depicted genomic interval are shown with arrows at boundaries (Patients 1, 2, 8, 9, and 10). Bottom panel. Consensus duplicated region of the 13 patients is depicted in the blue horizontal bar in a 405 kb window. Thirteen RefSeq genes are located in this region.

For DECIPHER entries with available parental information, 80% (n= 8) of observed microduplications were *de novo* and only one was inherited from an affected family member. The majority of (70%, 7/10) patients did not carry additional CNVs and none of the additional CNVs found in three patients covered a known disease locus or known disease genes. All 13 CNVs were highly heterogeneous in their size (average = 5.43 Mb; standard deviation = 4.41 Mb) and breakpoint distribution (encompassing from Chr19: 45.38 Mb to 59.09 Mb, Hg19) (Table 1).

To identify additional CNVs absent in ClinVar and/or DECIPHER databases, we screened the literature and retrieved three additional studies, including six patients with duplications at the 19q13.33 locus^11–13^. All of these patients suffered from seizures. Three patients carried CNVs of 1.22 Mb size whereas the remaining three duplications were >10 Mb. Patients affected by the large CNVs were, in addition to seizures, also affected by other NDDs including ID and dysmorphism. Interestingly, the two independent patients with the p.Val667Ile mutation on *GRIN2D* featured similar NDDs including developmental delay, dysmorphism, seizures and muscular hypotonia (Table 1).

Overall, the consensus duplicated region was determined to be located within the the coordinates 48,520,809 bp - 48,926,006 bp, with a final size of 405kb. This is consistent previous reports^11–13^. The consensus region overlapped 13 RefSeq genes (Figure 1B) that were further examined for brain expression, the presence of CNVs overlapping these genes in control cohorts and variation intolerance (Table 2). Four genes persisted above all available filters, namely: the caspase recruitment domain family member 8 (*CARD8)*; the chromosome 19 open reading frame 68 (*C19orf68)*; the KDEL endoplasmic reticulum protein retention receptor 1 (*KDELR1)*, and the glutamate ionotropic receptor NMDA type subunit 2D (*GRIN2D).* In our view these four genes represent the most promising candidates.

**Table 2.**
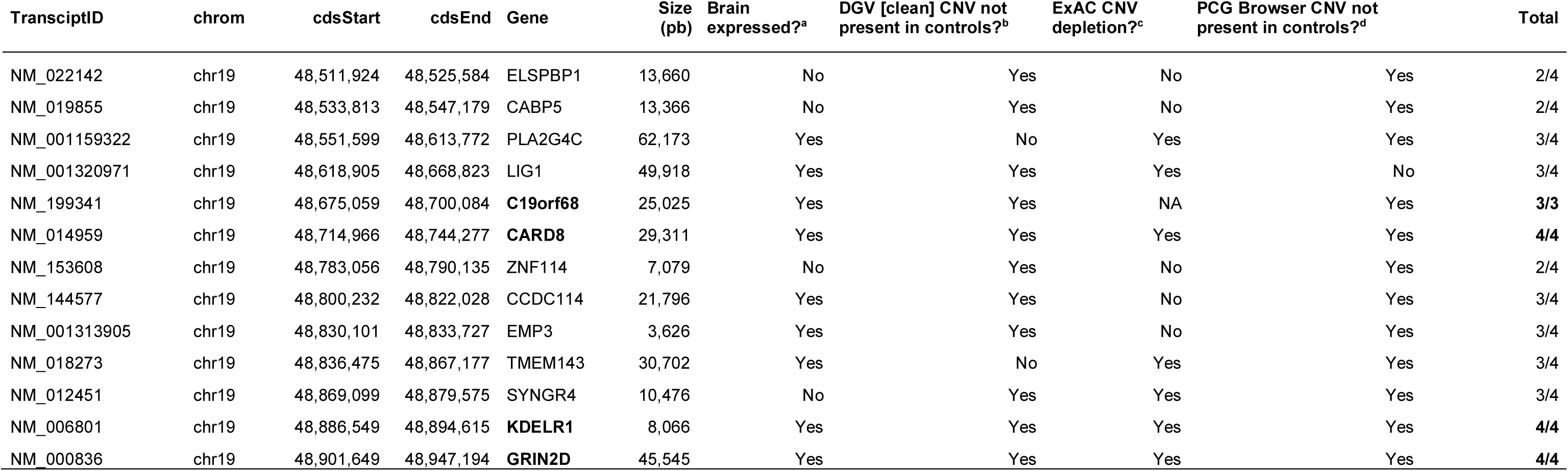
Consensus region gene annotation and candidate gene filtering.

We also searched for non-coding brain-specific functional elements within the consensus region. A total of 291 were found overlapping 9.87% of the consensus region (40,019 bp). Within the consensus regions of the duplications, the density of non-coding elements was not significantly higher than outside of chromosome 19.

## Discussion

Here we report on 13 patients with duplications at a potential novel disease locus within 19q13.33. Several lines of evidence support the hypothesis that duplications at this locus are associated with NDDs: I) Duplications at this locus are virtually absent in healthy individuals from the general population^8^; ll) All of the identified duplications with parental information arose *de novo* with the exception of patient 5 which according to DECIPHER was inherited from an affected parent with similar phenotype (DECIPHER entry 257554); III) None of the patients carried additional likely pathogenic CNVs and IV) all duplications covered multiple plausible disease candidate genes.

The NDDs observed in the 13 patients were characterized by dysmorphism as the most prominent feature, followed by intellectual disability and seizures (Table 1). Our observations are in agreement with previous reports^11–13^. Although, Wang *et al*, 2015^12^ focused exclusively on seizures and we cannot rule out that other NDDs were actually present in those patients. Similarly, we acknowledge that developmental delay, behavioral problems and learning difficulties may be subject to inter-observer variability to some extent. In this regard, future clinical studies of 19q13.33 duplication carriers need to be conducted to draw detailed and robust genotype-phenotype conclusions. Since previous reports from the literature were based on low resolution cytogenetic methods^13^, identification of the underlying disease gene was not possible. Here we show that by integration of multiple CNV datasets from public repositories we are able to narrow down the disease associated genomic sequence to a few candidate genes at 19q13.33 locus (Figure 1).

Our included datasets do not allow estimation of 19q13.33 duplication frequency. However, absence of 19q13.33 duplications in CNV databases of the general population and the presence of only a few variant carrying patients in diagnostic CNV databases with heterogeneous breakpoints indicate that 19q13.33 duplications are extremely rare (Table 2).

All 13 of the identified patient CNVs shared a genomic interval of 405 kb, which includes four genes with genetic, population and biological support of disease association. These included *CARD8*, *C19orf68*, *KDELR1* and *GRIN2D.* For *CARD8, C19orf68* and *KDELR1* no association with NDDs has been reported in the literature to date. Although we cannot rule out that brain-specific non-coding elements at 19q13.33 could be involved in the development of NDDs, *GRIN2D* represents a plausible candidate gene for association with NDDs. *GRIN2D*, encoding the NMDA receptor subunit GluN2D, is highly expressed prenatally and after birth before progressively declining through adulthood^14^. It is possible that *GRIN2D* microduplications may predispose to disease susceptibility in a dose dependent manner by enhancing GluN2D expression during development, thereby influencing the NMDA receptor composition, which might provoke changes in neuronal networks thus contributing to hyperexcitability and neurological diseases^15^. Besides CNVs, the *GRIN2D* gene is also depleted due to negative selection for missense and truncating variants in the general population, supporting the *GRIN2D* association with disease^8^. In agreement, two recently identified *GRIN2D* single nucleotide variants also lead functionally to a gain-of-function in two patients with similar outcomes^2^ (Table 1). Beyond the potential diagnostic relevance, our identification of *GRIN2D* as a possible new NDD gene has a potential clinical application, since memantine, a low-affinity therapeutic NMDA channel blocker, selectively blocks extrasynaptic NMDA receptors that are likely to contain GluN2C/2D subunits^16^. This might especially be relevant for patients with gain-of-function mutations or microduplications^2^.

## Acknowledgements

We thank all the clinicians, patients and their families. This study makes use of data generated by the DECIPHER community. A full list of centers who contributed to the generation of the data is available from http://decipher.sanger.ac.uk and via email from. Funding for the project was provided by the Wellcome Trust.

